# The astrocyte clock controls circadian perineuronal net remodeling, synapse strength and learning behavior

**DOI:** 10.64898/2026.04.04.716486

**Authors:** Philip C. Smith, Elsa Quillin, Katheryn B. Lefton, Celia A. McKee, Brendan Dang, Thomas Papouin, Erik Musiek

**Affiliations:** Departments of Anesthesiology, Washington University School of Medicine, St. Louis, MO, 63110; Departments of Neurology, Washington University School of Medicine, St. Louis, MO, 63110; Departments of Neuroscience, Washington University School of Medicine, St. Louis, MO, 63110; Center on Biological Rhythms and Sleep, Washington University School of Medicine, St. Louis, MO, 63110

**Keywords:** astrocyte, circadian, Bmal1, peri-neuronal nets, hippocampus, plasticity, LTP

## Abstract

The circadian clock controls a vast array of cellular and organismal functions, from the molecular scale to behavior. While each cell is regimented by a cell-autonomous clock, few studies in the brain have dissected the circuit and behavioral contributions of cell-specific clocks. Relatedly, astrocytes are now known to play key roles in regulating synaptic function, circuit activity and behavior, but whether these functions are guided by astrocyte-autonomous clocks is unknown. Here, we report that post-natal deletion of the critical circadian clock gene *Bmal1* in astrocytes, which abrogates core clock function in a cell type specific manner, induced expression of genes related to extracellular matrix (ECM) production, maintenance, and remodeling. Circadian variations have been shown in a specific ECM structure, perineuronal nets (PNNs), which are implicated in synaptic function and plasticity. In astrocyte-specific *Bmal1* knockouts, hippocampal PNN abundance was decreased, and the circadian rhythm of these structures was also abolished. In line with evidence implicating PNNs, and the ECM in general, in synaptic function and plasticity, we found that astrocyte-specific *Bmal1* KO mice had increased synaptic strength but blunted long term potentiation (LTP), as well as impaired learning and memory performance in a novel object recognition task. Taken together, these findings suggest that the astrocyte circadian clock regulates circadian rhythms in perineuronal net abundance as well as synaptic plasticity and behavioral learning and memory.

## Introduction

Circadian rhythms are endogenous biological rhythms entrained by zeitgebers (environmental cues, including light), and have been shown to be important for myriad biological processes. In addition to their best-known function as regulators of the sleep-wake cycle, circadian rhythms have been shown to have effects on metabolism, endocrine processes, cellular stress response, and many other core biological functions. The circadian clock has also been implicated in the control of neuronal function in the setting of learning and memory^1,2,3^. Multiple studies have shown a relationship between circadian rhythms and formation and consolidation of associative learning and declarative memories (for review see^4^). Consistently, hippocampal synaptic plasticity varies over the circadian cycle, strengthening during the active period of mice, and can be disrupted by genetic deletion of circadian clock genes in neurons^1,5,6,7^. However, cellular-molecular mechanisms of how circadian rhythms impact learning and memory remain a gap in our knowledge.

Anatomically, the central circadian clock is located in the Suprachiasmatic Nucleus (SCN) of the hypothalamus. While the SCN synchronizes circadian function to the light-dark cycle, remarkably, multiple brain cell types outside of the SCN have been shown to have their own endogenous circadian rhythms. In particular, glial cells such as astrocytes show daily rhythmicity in many brain areas^1,7,8^. Like in neurons, these cell-autonomous circadian rhythms are governed by the core circadian clock, which is dependent on the function of BMAL1 (Brain and muscle arnt-like, or Arntl), the key transcription factor of the circadian clock^9,10^. BMAL1 binds with either CLOCK or NPAS2 to form a nuclear heterodimer which drives circadian transcription of many genes, including *Per* (*Per1* and *Per2*), *Cry* (*Cry1* and *Cry2*), and REV-ERB (*Nr1d1* and *Nr1d2*). These negative-limb clock proteins then provide negative feedback regulation of BMAL1/CLOCK on a 24-hour cycle, forming the transcriptional basis of the circadian rhythm. Additionally, recent studies have shown BMAL1 dysfunction is implicated in a variety of neuropsychiatric^11,12^, endocrine/metabolism^13,14^, cardiovascular^15,16^, oncogenic^17,18,19^, and neurodegenerative disorders^12,20,21^. While the deletion of *Bmal1* has been used to probe the effects of cell-type-specific clock disruption, little is known at this time about the roles of the astrocyte-specific clock in brain functions or disorders.

Astrocytes have emerged as powerful regulators of synaptic function and plasticity, network activity, and behavior^22^. Astrocytes exhibit robust circadian rhythms^23^, especially in the SCN, where they regulate circadian behaviors^24,25,26,27^. In the hippocampus, circadian rhythms in astrocyte morphology and function are associated with circadian rhythms of synaptic plasticity and learning and memory^7^. Additionally, previous studies, including ours, have shown that the astrocyte-specific deletion of *Bmal1* impacts astrocyte activation, proteostasis, glutamate uptake, and the secretion of ATP, a well-studied astrocyte-derived transmitter^24,28,29,30,31^. Yet, the role of astrocyte clocks, and BMAL1 in particular, in the regulation of hippocampal learning at the synapse, circuit, and behavioral scales remains largely unknown and its mechanisms are not described.

Interestingly, an emerging mechanism of synaptic plasticity is through the regulation of the extracellular matrix (ECM)^32^. The ECM is the noncellular scaffolding component present within all tissues and organs, including the brain. It provides crucial chemical and mechanical input to cell behavior. (For review see^33^). Growing evidence suggests that the extracellular matrix (ECM) is regulated and remodeled by glial cells^34^ on a circadian basis^32,35,36^ (For review see^37^).

Perineuronal nets (PNNs) are specialized ECM structures^38^ that envelop the cell bodies and proximal dendrites of certain neurons in the brain, especially parvalbumin-positive interneurons^39^, and have been shown to regulate synaptic functions and plasticity^40^. PNNs have been implicated in brain development, particularly in the closure of the critical period for brain plasticity^6,41,42^. PNNs were classically conceptualized as stable structures after such a critical period closed in early childhood. However, recent research suggests that PNNs are dynamic and contribute to the regulation of neuronal and synaptic activity^32,35,36^. In particular, PNNs have been shown to dynamically respond during learning^43^, suggesting they may be important for regulation of the formation of new synapses^44^. PNNs have also been shown to be regenerated and modified on a diurnal time scale with a circadian rhythm^35,36^. Notably, PNN components are cleaved by matrix proteases that are diurnally expressed in the mouse cortex, such as cathepsin-S and MMP9^36,45,46^. Moreover, these rhythms coincide with dendritic spine density rhythms^36^. PNN rhythms are disrupted by sleep deprivation^36^, suggesting a potential structural connection between synaptic regulation important for brain plasticity and the circadian cycle.

Importantly, PNN content and structure are orchestrated by astrocytes and microglia, but PNN rhythms persist after microglial depletion^32^. Together, these data suggest a causal relationship between astrocytes, circadian cycles, ECM/PNNs, and synapse function.

In this study, we sought to investigate whether circadian rhythms in astrocytes modulate hippocampal synaptic plasticity, hippocampal learning, and the ECM. We disrupted the cell-autonomous circadian clock in astrocytes via *Bmal1* deletion, and found a cluster of changes in gene expression relating to ECM. Accordingly, astrocyte *Bmal1* knockout mice showed reduction in the number of hippocampal PNNs, as well as synaptic electrophysiological changes, including blunting of hippocampal LTP, and decreased performance in a hippocampal-dependent learning and memory task. Taken together, these findings suggest that the circadian clock in astrocytes regulates the extracellular matrix and perineuronal nets, and plays a key role in controlling the molecular and cellular processes underlying behavioral learning.

## Methods

### Animal care and use

To investigate the impact of Bmal1 in astrocytes, we employed a line of transgenic mice with inducible Cre-dependent selective deletion of *Bmal1* in astrocytes (*Aldh1l1*-Cre^ERT2^;*Bmal1*^f/f^), which we have previously characterized^28,47,29^. All animal experiments were approved by the Washington University IACUC and were conducted in accordance with AALAC guidelines and under the supervision of the Washington University Department of Comparative Medicine. *Aldh1L1-Cre^ERT^*^2^ and *Bmal1^fl/fl^* mice were obtained from The Jackson Laboratory (Bar Harbor, ME) and were bred at Washington University. All cohorts of mice were mixed sex and consisted of Cre^+^ and Cre^−^ littermates from several breeding cages; mixed-sex cohorts were used as no sex differences have been observed in this model. All mice were maintained on a C57Bl/6 background and housed under 12-h light/12-h dark conditions. All mice (Cre^−^ and Cre^+^) were given tamoxifen (Millipore Sigma, Billerica, MA, dissolved in corn oil, 2 mg/mouse/day for 5 d) by oral gavage at 2 mo of age to induce *Bmal1* astrocyte-specific deletion in Cre^+^ mice. All mice were group housed with food and water available *ad libitum.* At the conclusion of each experiment, mice were deeply anesthetized with intraperitoneal pentobarbital (150mg/kg), then perfused with ice-cold Dulbecco’s modified PBS (DPBS) containing 3 g/l heparin. Brain tissue was then harvested for sectioning and gene analysis. One hemisphere was rapidly dissected and flash frozen, while the other was drop-fixed for 24 hours in 5% paraformaldehyde for 24 h (4 °C), then cryoprotected with 30% sucrose in PBS (w/v, 4 °C) until sectioning.

### RNA Sequencing and Gene Analysis

We re-analyzed existing RNA sequencing data from bulk cortex tissue from astrocyte-specific Bmal1 knockout (Bmal1^KO^) mice, as described in our previous work^47^. We employed a permissive cutoff of genes upregulated in Cre^+^ (astrocyte Bmal1^KO^) mice with an uncorrected P<0.05, then analyzed this list for Gene Ontology using the Database for Annotation, Visualization, and Integrated Discovery (DAVID, NIH, Bethesda, MD)^48^. The list of differentially-expressed genes used for DAVID analysis is available in Supplemental Table 1.

### WFA-Parvalbumin Co-Staining for Perineuronal Nets

Fixed brain tissue was frozen and cut into 40 µm thickness coronal sections using a sliding microtome (Leica, Deer Park, IL). Using stereotactic sections described previously^20^, slices were identified corresponding to the CA3 and CA1 regions of the hippocampus. Slices were washed for 3 iterations of 10 minutes each in phosphate-buffered saline (PBS) (Corning, Corning, NY) and then permeabilized for 1 hour in 0.3% Triton with PBS (Millipore Sigma). Slices were washed as before and blocked for 1 hour in 10% goat serum (Abcam, Cambridge, MA). They were then incubated for 36 hours at 4 degrees Celsius, in a solution of PBS with 3% goat serum, 0.3% Triton, 1:1000 WFA-Biotin antibody (Vector Laboratories, Newark, CA) and 1:1000 mouse anti-parvalbumin antibody (Sigma-Aldrich, St. Louis, MO). After incubation, the slices were washed for 4 iterations of 10 minutes each as above. Slices were then incubated 2 hours at room temperature in 0.3% Triton, 3% goat serum PBS with 1:1000 dilution of goat-anti-mouse 568 (Life Technologies, Carlsbad, CA) and 1:1000 dilution of streptavidin 488 (Life Technologies). Finally, slices were washed 5 times in PBS and mounted with Prolong Gold Mounting Medium (Thermo-Fisher, Waltham, MA). Slices were visualized on a Keyence BZX-800 microscope (Keyence, Itasca, IL). PNN density measures were obtained by examining two 40µm sections per animal and counting all complete PNNs or PV neurons observed in the entire hippocampal region, as applicable, and dividing by the total visualized area to obtain density metrics.

### Electrophysiology

Electrophysiology experiments were carried out in acute hippocampal coronal slices (350 µm) obtained with a Leica VT1200s vibratome from adult male and female mice (P90-120) as previously described^49,50,51^. After recovery (30min at 33°C and 45min at room temperature), slices were transferred to the recording chamber of a SliceScope Pro 6000 system (Scientifica, Miami, FL), where they were perfused with artificial cerebrospinal fluid (aCSF) saturated with 95%O2/5%CO2 at a flow rate of ∼1mL/min. The aCSF was maintained at 33°C (TC-344C Dual Channel Temperature Controller, Warner Instruments, Holliston, MA). The aCSF composition for slicing and recovery was (in mM) 125 NaCl, 3.2KCl, 1.25NaH2PO4, 26 NaHCO3, 1 CaCl2, 2 MgCl2, and 10 glucose (pH 7.3, 290-300 mOsm.L-1). The aCSF composition for recording was (in mM) 125 NaCl, 3.4KCl, 1.25NaH2PO4, 26 NaHCO3, 2 CaCl2, 1.3 MgCl2 and 10 glucose (pH 7.3, 290-300 mOsm.L-1). Schaffer collaterals were electrically stimulated with a concentric tungsten electrode placed in the *stratum radiatum* of CA1, using paired stimulations (100 μs pulse, 200 ms apart) continuously delivered at 0.05 Hz. Evoked field excitatory post-synaptic potentials (fEPSPs) were recorded using a glass electrode (2-5 MΩ) filled with recording aCSF and placed in the *stratum radiatum* near the isosbestic point. Stimulation intensity (<150 μA) was set as needed to evoke a synaptic response without population spikes within the slope of the fEPSPs. Experiments were performed at 33° C in the presence of the GABAA receptor blocker picrotoxin (50μM, Tocris, Minneapolis, MN). Data were acquired with a Multiclamp 700B amplifier (Molecular Devices, San Jose, CA) through a Digidata 1450A, sampled at 20 kHz, filtered at 10 kHz, and analyzed using pClamp11.0 software (Molecular Devices). Average traces were taken from 10min epochs before and after drug application at times indicated by numbers in parenthesis on time-course graphs. The paired-pulse ratio was measured as the ratio of the slope of the second fEPSP over the slope of the first fEPSP. Ratios greater than 1 (paired-pulse facilitation, PPF) are typical at CA3-CA1 synapses and reflect an enhanced glutamate release during the second synaptic response. This short-term potentiation is due to the incomplete Ca^2+^ buffering in pre-synaptic terminals during the short inter-pulse interval, leading to an additive effect on pre-synaptic free Ca^2+^ levels and an exaggerated vesicular release in response to the second pulse. For LTP experiments, stimulation intensity was set to 35% of that triggering population spikes. After a stable baseline of 20 min, LTP was induced by applying a High Frequency Stimulation (HFS) protocol consisting of a three 100 Hz trains of stimulations delivered for 1 s, 20 s apart.

### Novel Object Recognition Assay

The novel object recognition task was run in a quadruple open-field setup described previously^52^ over three days, at ZT6, under 150lux. All mice were handled by the same experimenter (male). There was no prior habituation to the human handler. Day 1 consisted of an acclimation trial wherein mice were free to explore the empty arena for 10min, after a 15min acclimation to the experimental room, then immediately returned to their home cage and to the vivarium. Day 2 consisted of two successive acquisition sessions (30min apart), wherein mice were reintroduced in the same chamber with two identical objects were present and were free to explore the objects for 10min. During the inter-session interval, mice were returned to their home cage in the experimental room. Day 3 consisted of a probe trial wherein one of the two objects was replaced with a new (never-before-encountered) object. The location of the new object was balanced across cohorts to counter-balance object place preferences from individual mice. The objects were similar in proportions but distinct in material (plastic vs. ceramic, height, texture, color and shape). Two to four littermates of same sex were run simultaneously. A three-point (nose, center of mass and tail) mouse tracking and object exploration were performed online using Smart3 software (Harvard Apparatus, Holliston, MA) and analyzed off-line. Object exploration was defined as a direct contact with the object or directed probing within 3.5 cm of the object. The discrimination index (DI) was measured as t_novel_ – t_familiar_ / (t_novel_ + t_familiar_). The novel object place preference index computed during the acquisition phase represents the preference (or lack thereof) for the object position where the novel object is later introduced.

## Results

### Astrocyte Bmal1 regulates ECM-related gene expression in a circadian-dependent manner

To investigate the impact of BMAL1 in astrocytes, we employed a line of transgenic mice with inducible Cre-dependent selective deletion of *Bmal1* in astrocytes (*Aldh1l1*-Cre^ERT2^;*Bmal1*^f/f^), which we have previously characterized^28,29,47^. This mouse line presents the advantage of being able to isolate the impact of astrocyte circadian rhythm on downstream processes without the confounding factors presented by deletion of circadian regulation in other cell types. We analyzed existing bulk RNA sequencing data from cortical tissue from these animals^47^, employing a permissive cutoff of genes upregulated in Cre^+^ (astrocyte Bmal1 knockout (Bmal1^KO^)) mice with an uncorrected P<0.05 (Table S1). We then interrogated this list for gene ontology using the DAVID database^48^ (Fig. 1A). Our analysis revealed several enriched pathways, including a cluster of genes involved in extracellular matrix (ECM) production and remodeling (Fig. 1B). We focused on these pathways due to previous studies implicating ECM in regulation of synaptic plasticity^53,54^. The gene with the most prominent degree of change was *Mmp14*, a matrix metalloproteinase that regulates ECM degradation^55^, which was upregulated by 2.4-fold in the transcriptome of Bmal1^KO^ mice at ZT 6 relative to their Cre-negative littermate controls. To understand if the expression of these transcripts is regulated in a circadian fashion, we examined our circadian RNAseq database obtained from mouse cortex (created using astrocyte-specific translating ribosome affinity purification^56^). We found within the cluster of ten upregulated transcripts in Bmal1^KO^ mice, four of those showed strong circadian rhythmicity in astrocytes, including *Mmp14, Palld, Emp2, and Bag3*, as determined using RAIN algorithm^57^ (Fig. 1C). To examine the effect of *Bmal1* deletion on these genes at different times of day and across models, we examined expression in cortex from brain-specific *Bmal1* KO animals (*Nestin*-Cre;*Bmal1*^f/f^ mice, as described previously^28^). These four transcripts were also upregulated significantly at ZT12, while *Mmp14, Emp2,* and *Bag3* were also significantly increased at ZT0 (Fig. 1D) (p<0.05 by two-way ANOVA with post-hoc Sidak’s multiple comparison’s test). Thus, the core circadian clock in astrocytes regulates expression of multiple genes involved in ECM homeostasis, and these genes are generally upregulated following *Bmal1* deletion.

**Figure 1:**
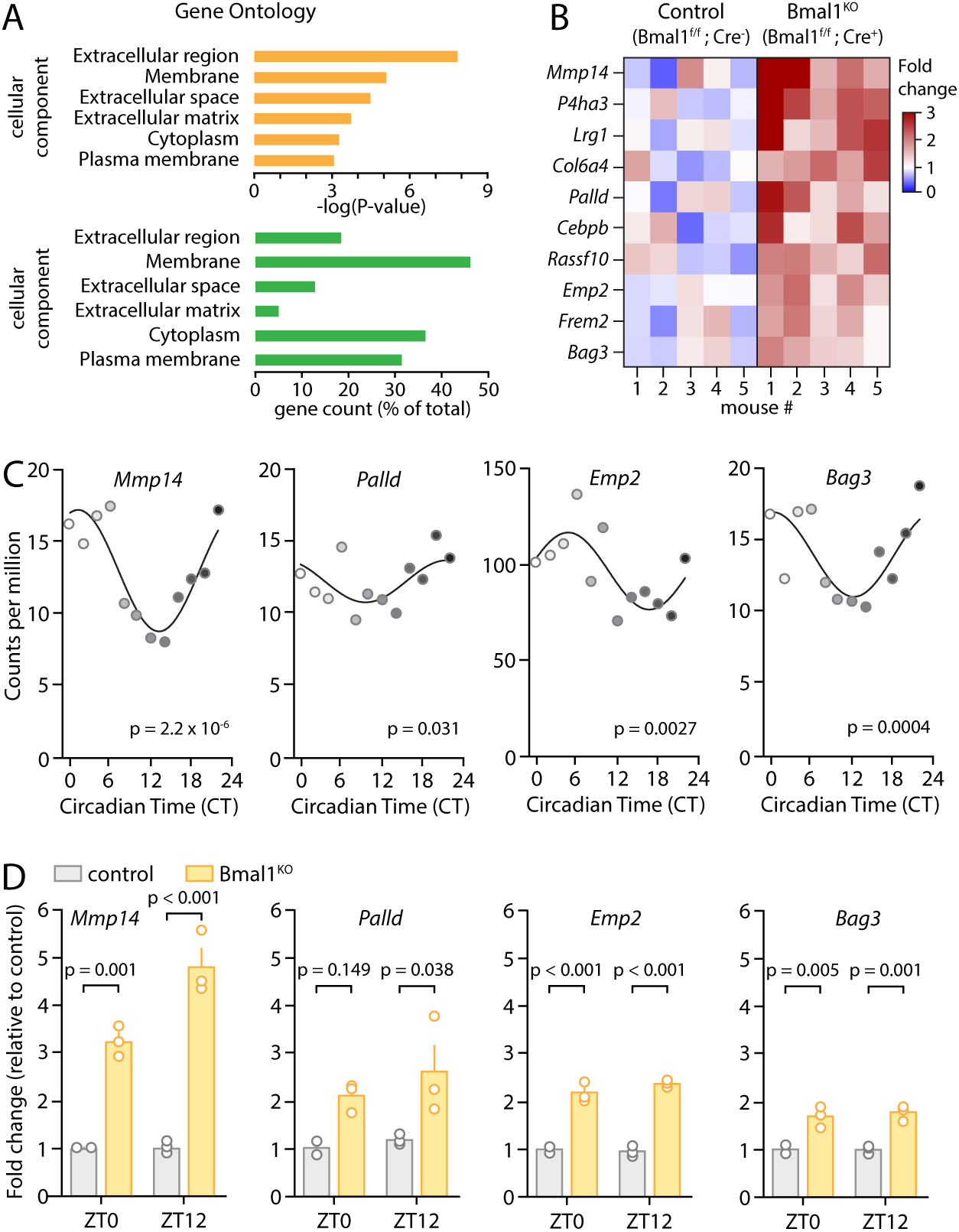
Astrocytic Bmal1 deletion changes expression of extracellular matrix genes. (**A**) Pathway analysis of differentially expressed genes in cortex tissue from astrocyte-specific Bmal1 knockout mice (Bmal1^KO^). The Gene Ontology – Cellular Component (GO-CC) database was queried using the Database for Annotation, Visualization, and Integrated Discovery (DAVID)^48^. The -log of the P value for enrichment is shown. Extracellular matrix (ECM) was among the most enriched cellular components (p=1.17*10^-5^). (**B**) Heatmap of expression of ECM-related genes for individual control and Bmal1^KO^ animal (n=5). (**C**) Circadian gene expression of *Mmp14, Palld, Emp2*, and *Bag3* in wild type astrocytes from a circadian gene atlas derived from previous studies^56^. P values indicate rhythmicity based on analysis with RAIN^57^. (**D**) *Mmp14, Emp2*, and *Bag3* are significantly upregulated in cortex tissue from brain-specific Bmal1^KO^ animals (*Nestin*-Cre;*Bmal1*^f/f^) at ZT0 and ZT12, with *Palld* upregulated at ZT12 only. Two-way ANOVA with post-hoc Sidak’s multiple comparisons test was used.

### Astrocyte BMAL1 regulates circadian rhythms in PNN content

Having showed that *Bmal1* deletion in astrocytes affects the expression of genes related to ECM maintenance, we next sought to investigate the effect of astrocytic *Bmal1* KO on ECM structures. The best studied component of the ECM in relation to neuronal activity and synaptic function are perineuronal nets (PNNs), which are extracellular collections of proteoglycans that specifically form around GABAergic parvalbumin-positive (PV^+^) interneurons^44,58,59,60^. PNNs gate inhibitory PV^+^ interneuron (PV-INs) activity and GABAergic transmission in the hippocampus, with the loss of PNNs causing GABA-mediated suppression of LTP^59,61^. Moreover, the number of PNNs in the hippocampus varies in a circadian manner^32,35,36^, with a trough around ZT6 and peak around ZT18. We thus next examined PNN abundance, and potential circadian rhythmicity, in astrocyte-specific Bmal1^KO^ mice and their littermate controls at these times of day. We treated all mice with tamoxifen at 2 months old to delete astrocytic *Bmal1* in Cre^+^ mice, then harvested mice under constant darkness (DD) conditions (for 24 hours prior to and during harvest) at 4 months of age. We performed co-staining of PNNs (*Wisteria Floribundii Aglutinin* (WFA))^62^ and PV^+^ neurons and identified PNNs that colocalized with PV^+^ neurons throughout the hippocampus. We performed counts of all PNNs in the hippocampus for animals harvested at ZT6 and ZT18 (Fig. 2A-B) and analyzed data by 2-way ANOVA. The main effect of genotype on PNN density was highly significant (p<0.0001), as PNN density was significantly decreased across the hippocampus in Bmal1^KO^ mice at both time points (Fig. 2B). The main effect of time was not significant. However, there was a strong trend toward a positive interaction between genotype and time (p=0.056), with the density of PNNs trending toward increase at ZT18 over ZT6 in control mice, but not in Bmal1^KO^ mice. At ZT6, we found a decrease in Bmal1^KO^ mice in both raw PNN count across the CA1 hippocampus and percentage of the hippocampal area staining positive for PNNs (p=0.013, unpaired Student’s *t*-test) (Fig 2D). The change in PNN density occurred despite no change in the number of PV^+^ neurons (p=0.71, unpaired Student’s *t*-test) (Fig. 2C). These results demonstrate that the astrocyte circadian clock regulates daily rhythms in PNN abundance in the hippocampus, and that astrocyte *Bmal1* KO mice have decreased hippocampal PNN content.

**Figure 2:**
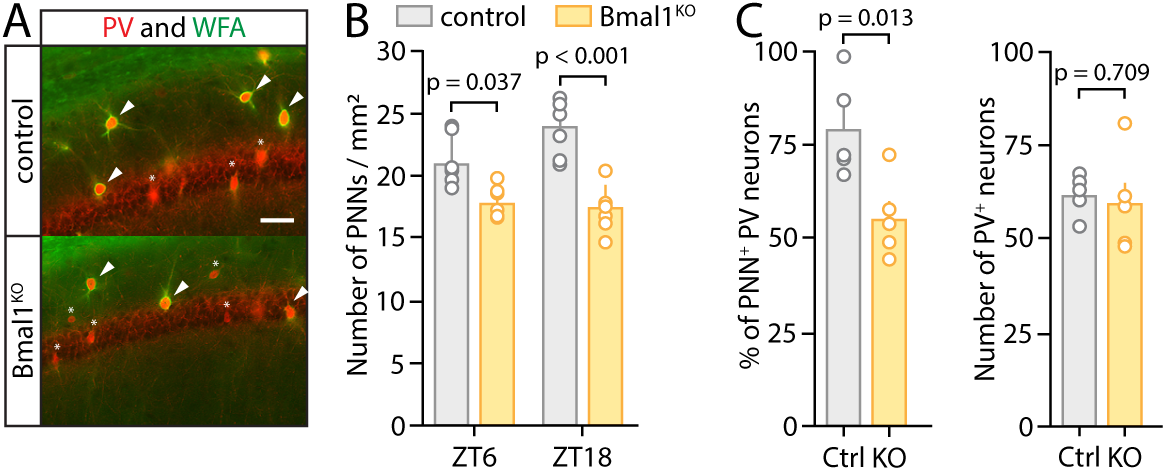
Astrocytic Bmal1 deletion changes expression of perineuronal nets in a circadian-dependent manner. (**A**) Representative confocal images of hippocampal sections from control and astrocyte-specific Bmal1^KO^ animals, taken at ZT 6, and co-stained for parvalbumin (PV, red) and PNNs (WFA, green). (**B**) Quantification of PNNs density in the hippocampal CA1 region (n= 6/group). Interaction between time and genotype analyzed by 2-way ANOVA with post-hoc Tukey’s multiple comparisons test. Main effect of genotype (p<0.0001) was significant, as well as a trend toward main effect of interaction between time and genotype (p=0.056). (**C**) *Left*, quantification of the percentage of parvalbumin-positive (PV^+^) neurons which are surrounded by a PNN, in sections from animals euthanized at ZT 6 (each circle indicates data from one mouse and mean+SEM is shown, p=0.013, unpaired Student’s t-test). *Right*, quantification of number of PV^+^ neurons in the CA1 region with no differences between control and Bmal1^KO^ sections (p=0.71, unpaired Student’s t-test). 2-way ANOVA with post-hoc Tukey’s multiple comparisons test (B) and unpaired Student’s t-test (C) (D) were used.

### Astrocyte *Bmal1* deletion augments synaptic strength and blunts plasticity

Given the role of both astrocytes and PNNs in governing synaptic function, we next examined the effect of the astrocyte-specific deletion of *Bmal1* on hippocampal synaptic function and plasticity, focusing on the ZT6 timepoint. To assess synaptic function at CA3-CA1 synapses, we performed extracellular recordings of AMPAR-mediated field excitatory postsynaptic potentials (fEPSPs) in the *stratum radiatum* of acute hippocampal slices taken from adult Bmal1^KO^ and control mice. We first assessed baseline synaptic strength by measuring the fEPSP slope in response to increasing intensities of stimulation (20-120 µA). This input–output relationship revealed an exaggerated synaptic strength in astrocyte Bmal1^KO^ mice compared with control animals, synonymous of an overall enhancement in glutamatergic transmission in Bmal1^KO^ mice (Fig. 3A). This coincided with increased excitability of Bmal1^KO^ slices overall (e.g., pop spikes, not shown). Measurement of the paired-pulse facilitation (Methods) in a subset of the same animals, over a range of interstimulus intervals (25, 50, 100, 200 and 500 ms), however, revealed no significant difference between Bmal1^KO^ and control mice (p>0.05, multiple comparison ANOVA), indicative that the probability of release at CA3-CA1 synapses was not affected by the astrocyte *Bmal1* deletion (Fig. 3B). Next, we examined hippocampal long-term synaptic plasticity in control and KO animals, using a classic tetanic LTP-inducing protocol (HFS: 3 x 1s, 100Hz, Methods) delivered after a 20min baseline. In slices from control animals, such a protocol produced a long-lasting strengthening of synaptic transmission (153.0 ± 5.9%, n = 11 mice) (Fig. 3C), which was significantly reduced in the slices from Bmal1^KO^ mice (136.5 ± 4.0%, n = 10 mice; p = 0.0293, unpaired Student’s *t*-test). This was not due to an increased threshold for LTP induction, but rather to a population shift in the percent of potentiation (Fig. 3C). Together, the increased input-output relationship, reduced ability to potentiate, and intact PPF, suggest that synapses in Bmal1^KO^ slices are in a constitutively enhanced, near-saturated state.

**Figure 3:**
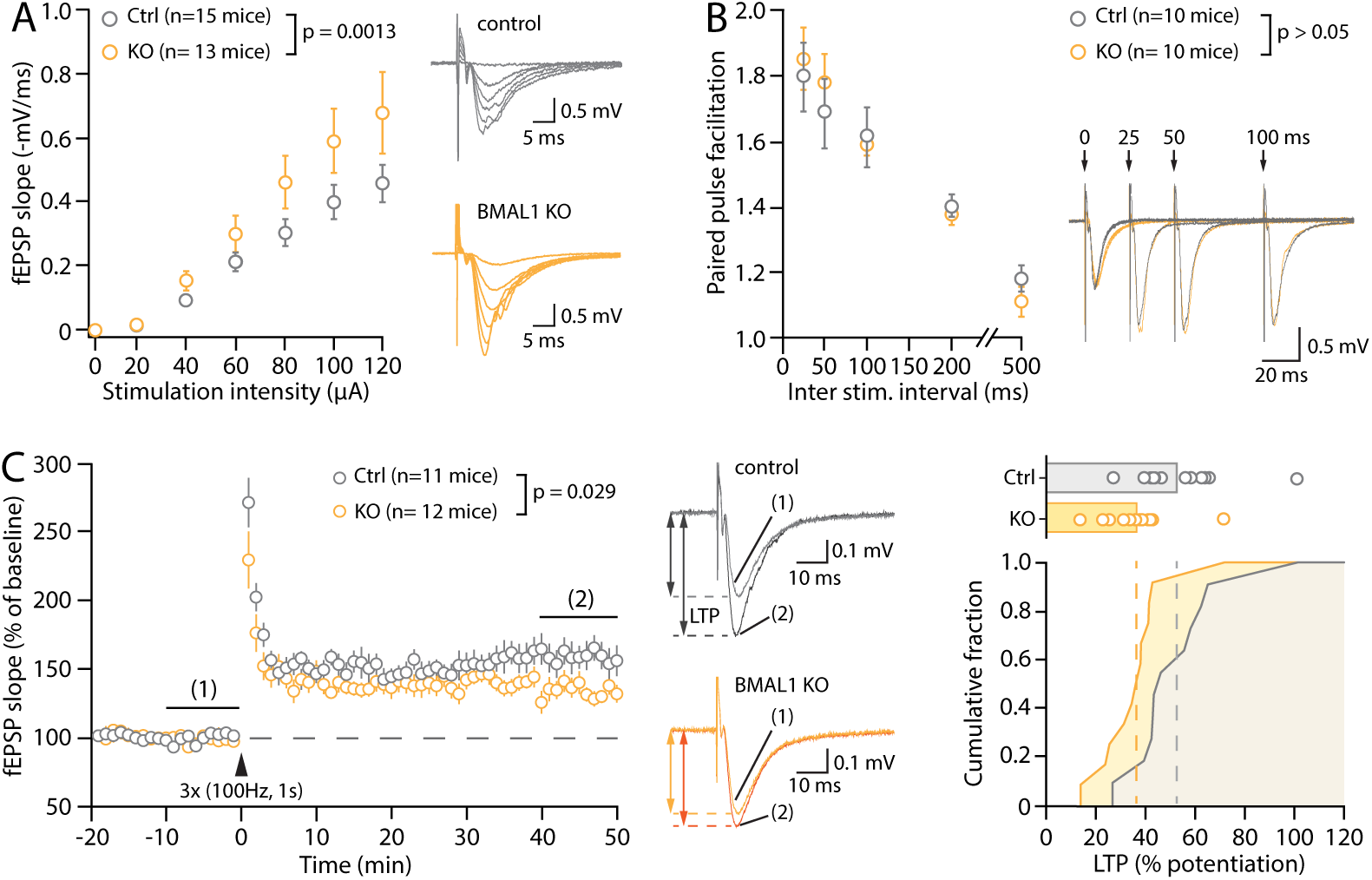
Effect of astrocyte-specific Bmal1 deletion in astrocytes on synaptic function. (**A**) Input-output relationship of synaptic fEPSPs recorded in the CA1 *stratum radiatum* of hippocampal slices from control and Bmal1^KO^ adult mice and example traces with overlaid responses at increasing intensities. The fEPSP slope/stimulation intensity relationship is greater in BMAL1^KO^ slices. (**B**) Paired-pulse facilitation of the second response when paired stimuli are delivered at the indicated inter-stimulus interval (ISI), and illustrative traces. Note that for clarity the illustrative responses to 200ms and 500ms ISI are not shown. (**C**) *Left*, time-course of fEPSP slope over time in response to 3 tetanic trains (100Hz, 1s) delivered 20 seconds apart (indicated by black arrowhead). Each circle represents the average fEPSP slope of 3 consecutive sweeps (i.e. 1min). The illustrative traces show representative fEPSPs averaged from 30 sweeps in the baseline and LTP conditions (taken at the indicated times on the time-course). *Right*, cumulative distribution of the percent of synaptic potentiation for all individual experiments. Multiple comparison ANOVAs (A, B) and unpaired Student’s *t*-test (C) were used.

### Astrocyte *Bmal1* deletion impacts novel object recognition

To probe the consequences of our molecular and physiological findings on hippocampal learning, we employed the novel-object recognition task. This hippocampus-dependent behavior paradigm takes advantage of mice having an innate preference for novelty to indirectly probe long-term semantic learning and memory^63^. After being exposed to two identical objects in an open-field arena (under low lighting, see Methods), mice returned to the same arena 24hrs later, with one of the objects replaced by a novel one (Fig. 4A). Whereas control animals showed a clear preference for the novel object (control: 39.2 ± 5.2s vs. 19.0 ± 2.7s, n = 19 mice, p < 0.001, paired Student’s *t*-test, Fig. 4B), indicative of normal long-term object recognition, astrocyte Bmal1^KO^ animals showed no such preference (36.7 ± 3.4 vs. 30.7 ± 4.1s, n = 20, p = 0.172, paired Student’s *t*-test), resulting in a significantly reduced discrimination index (control: 0.32 ± 0.05, n = 19; KO: 0.13 ± 0.05, n = 20, p = 0.0159, unpaired Student’s *t*-test, Fig. 4C-D), with no alterations in total object exploration time (Fig. 4E) or distance traveled (Fig. 4F) during either the acquisition or probe phase. Together, these results indicate that *Bmal1* deletion in astrocytes, which reduced synaptic plasticity and decreases hippocampal PNN content at ZT6, also results in a significant learning and memory deficit.

**Figure 4:**
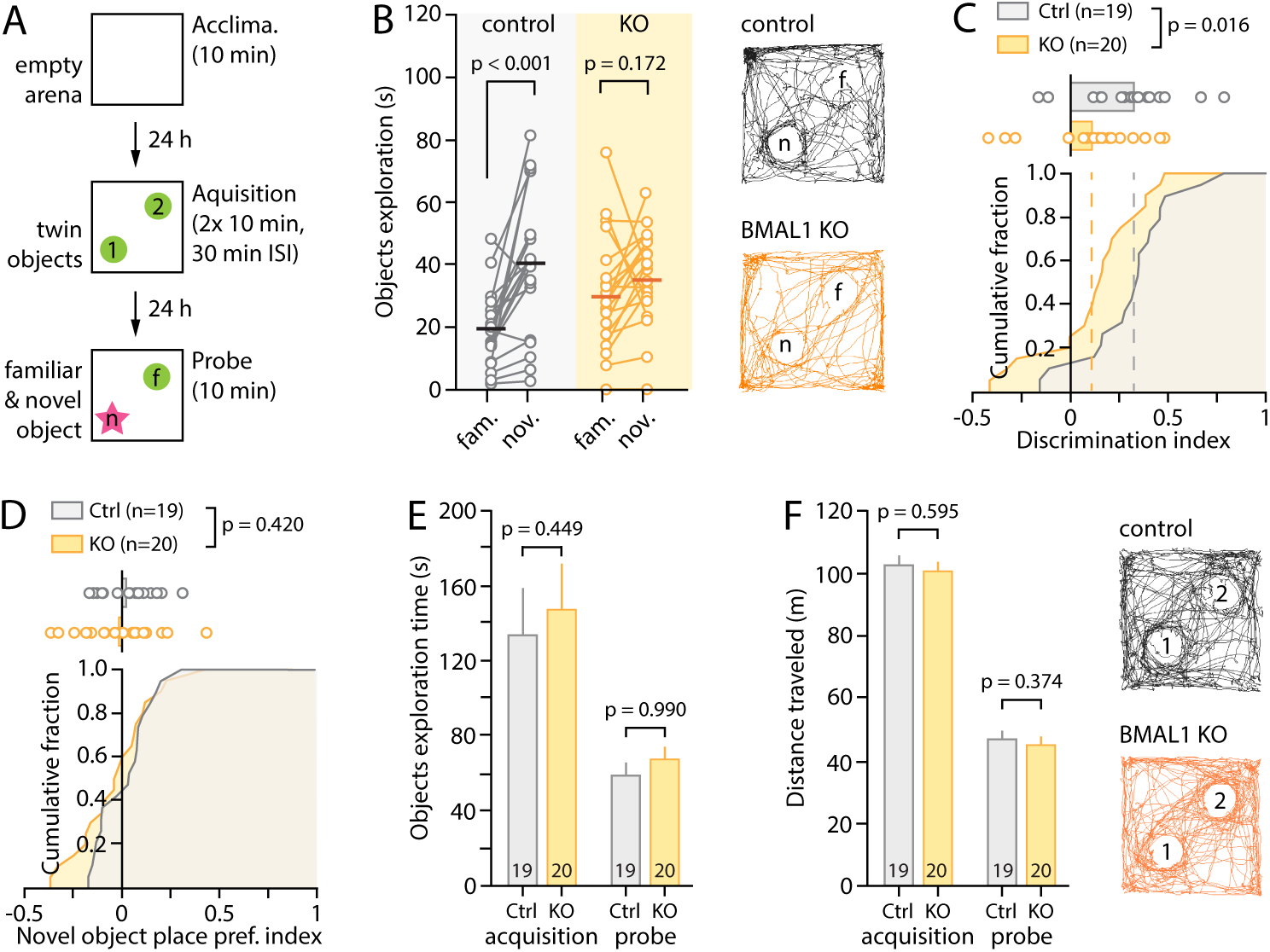
Bmal1 deletion in astrocytes impairs hippocampal learning. (**A**) Schematic illustrating the time-course of the novel object recognition task. (B) Object exploration times for individual mice during the probe phase, showing a clear preference of control animals for the novel object (P<0.001, paired Student’s t-test) that is abolished in Bmal1^KO^ mice (p=0.172). Illustrative activity traces are shown for the entire 10min of exploration during the probe phase. (C) Cumulative fraction and individual values of discrimination index computed for each mouse from the data shown in B (see Methods). The performance of the Bmal1^KO^ mice is markedly reduced compared to that of the controls (p=0.016, unpaired Student’s t-test). (**D**) Cumulative fraction and individual values of novel object place preference index computed from exploration times during the acquisition phase and showing that neither genotype had a biased preference for the spatial location where the novel object was placed on the following probe day. (**E-F**) Bar graphs showing the total object exploration times (both objects combined, E) and distance traveled (F) during the acquisition and probe day. Illustrative activity traces are shown for the entire 10min of exploration during the acquisition phase. Paired Student’s *t*-test (A) and unpaired Student’s *t*-tests (C-F) were used.

## Discussion

In this manuscript, we show that the master circadian regulator *Bmal1* in astrocytes impacts neuronal gene expression, anatomy, circuitry, and behavior related to learning and memory. Astrocytic *Bmal1* deletion changes the expression of multiple genes involved in extracellular matrix production, maintenance, and turnover, many of which are regulated in a circadian manner. At the protein level, we found that astrocytic *Bmal1* deletion reduces the expression of perineuronal nets in the hippocampus. At the level of neuronal circuitry, we show that *Bmal1* astrocytic deletion results in enhanced synaptic strength and a decrease in synaptic potentiation in a circuit key for learning and memory, consistent with PNN removal lifting a brake on plasticity, and in line with prior findings^61^. Consistently, at the level of behavior, we find that astrocytic *Bmal1* deletion results in decreased performance in hippocampal learning and memory. Taken together, these results show that the core clock gene BMAL1 in astrocytes controls a cascade of downstream effects on neuroanatomical structures and circuitry involved in learning and memory, as well as on behavioral tests of such memory activity. While neuronal circadian rhythms are known to regulate daily variation in hippocampal synaptic plasticity and learning and memory, our results provide evidence that the astrocyte circadian clock contributes to these processes, potentially through regulation of PNNs.

Previous work from our group and others has demonstrated key functions on BMAL1, and the circadian clock in general, in the regulation of astrocyte gene expression and reactivity state. Tissue-specific deletion of *Bmal1* can be used as a strategy to probe the effects of cell-type-specific clock disruption, though it should be noted that BMAL1 exerts some non-circadian effects^21^ which could result in off-target effects in this model. Astrocytes show robust circadian rhythms in clock gene expression in culture, and the astrocyte clock has been implicated in regulation of specific functions such as ATP and glutamate release and reuptake^30^. In vivo, astrocytes have an abundant circadian transcriptome^64^ and *Bmal1* deletion causes robust, cell-autonomous astrocyte reactivity phenotype with changes in gene expression and lysosomal function^28,29,47^. Glial clocks in drosophila have been thoroughly implicated in control of rhythmic behavior^65^. In the suprachiasmatic nucleus of mice, astrocyte circadian rhythms are known to be critical for maintenance of neurotransmitter rhythms and are critical for maintaining SCN function and whole-animal behavioral rhythms^24,25,26,27^. Astrocyte Bmal1 in hypothalamic nuclei has also been implicated in regulating energy expenditure and metabolism in mice^66,67,68,69^, while deletion in the nucleus accumbens impacts reward behavior^64^. Thus, there is some precedent for the astrocyte clock mediating neuronal function, though this had not been shown in the context of hippocampal plasticity or learning and memory. Indeed, synaptic plasticity and learning and memory do show robust circadian rhythmicity which can be abrogated by global or neuron-specific *Bmal1* deletion, but was explained by cell-autonomous events within neurons, including alterations in MAP kinase signaling, GSK3β function, or synaptic BMAL1 phosphorylation^1,2,70,71,72,73^. Our findings suggest an additional level of control featuring astrocyte clock-mediated regulation of PNNs, providing evidence of circadian cell non-autonomous regulation of synaptic plasticity in the hippocampus. It remains unknown how this astrocyte-mediated PNN mechanism interacts with neuron-intrinsic circadian processes to regulate learning and memory rhythms, and this is a promising area of future inquiry.

Beyond *Bmal1*, these studies present the possibility of a mechanism for astrocytic circadian rhythms impacting learning and memory through extracellular PNN regulation. Perineuronal nets have long been established as important for learning in the context of the critical period for brain development^74^, and recent research has shown that they may play a role in stabilizing synapses for potentiation^75,76,77^ (For review see^78^). The effects of *Bmal1* shown here at the level of gene expression, ECM protein structures, electrophysiological synaptic activity, and behavior suggest a mechanism involving PNN regulation, though our data does not prove a direct causal relationship and future work is needed to fully elucidate this mechanism. Other unidentified astrocyte genes and/or ECM structures may also contribute to these effects. Moreover, ECM regulation by microglia has been shown to impact synaptic plasticity^53,65,54^, so effects of microglial clocks, or indirect effects of astrocyte clock function on microglia, could be considered in future studies. However, our work here provides clear evidence that astrocyte Bmal1 deletion disrupts PNN expression and alters synaptic plasticity and behavior, laying the groundwork for further examination of the role of astrocyte circadian clocks in the control of ECM structures and learning and memory.

## Acknowledgements

This project was supported by NIH grants R01AG054517 (to EM) and R01MH127163-01 (to TP). Involvement of PS was supported by the Washington University Department of Anesthesiology Training Grant T32GM108539. Confocal microscopy was performed in part through the use of Washington University Center for Cellular Imaging (WUCCI) supported by Washington University School of Medicine, The Children’s Discovery Institute of Washington University and St. Louis Children’s Hospital (CDI-CORE-2015-505 and CDI-CORE-2019-813) and the Foundation for Barnes-Jewish Hospital (3770 and 4642).

---

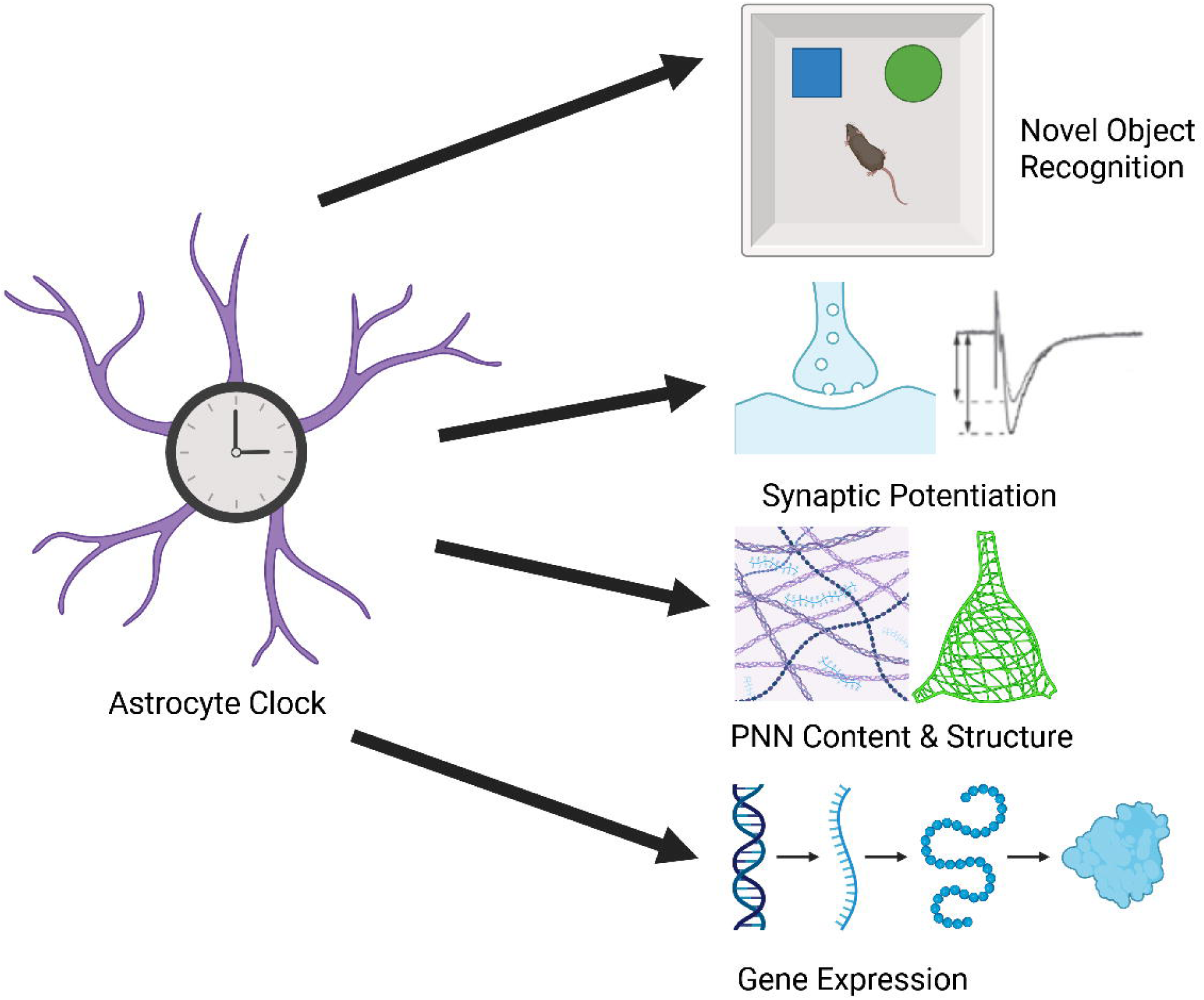

